# Choice between 1- and 2-furrow cytokinesis in *Caenorhabditis elegans* embryos with tripolar spindles

**DOI:** 10.1101/450478

**Authors:** Tomo Kondo, Akatsuki Kimura

**Author notes:** Correspondence to Akatsuki Kimura: Cell Architecture Laboratory, Structural Biology Center, National Institute of Genetics, Yata 1111, Mishima, Shizuoka 411-8540, Japan., ORCID: 0000-0003-4227-4811. Graduate School of Sciences and Technology for Innovation, Yamaguchi University, 1677-1 Yoshida, Yamaguchi City, Yamaguchi 753-8512, Japan (Tomo Kondo).

## Abstract

Excess numbers of centrosomes often lead to multipolar spindles, and thus probably to multipolar mitosis and aneuploidy. In *Caenorhabditis elegans*, approximately 70% of the paternal *emb-27^APC6^* mutant embryonic cells contained more than 2 centrosomes and formed multipolar spindles. However, only 30% of the cells with tripolar spindles formed 2 cytokinetic furrows. The rest formed 1 furrow, like normal cells. To investigate the mechanism how the cells avoided to form 2 cytokinetic furrows even with a tripolar spindle, we conducted live-cell imaging in *emb-27^APC6^* mutant cells. We found that the chromatids were aligned only on 2 of the 3 sides of the tripolar spindle, and the angle of the tripolar spindle relative to the long axis of the cell correlated with the number of cytokinetic furrow. Our numerical modeling showed that the combination of cell shape, cortical pulling forces, and heterogeneity of centrosome size determines whether cells with tripolar spindle form 1 or 2 cytokinetic furrows.

## Introduction

The centrosome is a major microtubule-organizing center in animal cells. Each centrosome contains a pair of centrioles, and they duplicate only once during a cell cycle. Therefore, the number of centrosomes in a cell is strictly regulated (Nigg and Holland, 2018). Normally dividing cells have 2 centrosomes that become the 2 poles of the bipolar mitotic spindle to segregate the sister chromatids into 2 daughter cells after mitosis. Through the microtubules elongating from the centrosomes, the centrosomes function as a hub to aggregate forces acting on the microtubules, such as cortical pulling forces, cytoplasmic pulling forces, or cortical pushing forces (Reinsch and Goncy, 1998; Grill and Hyman, 2005; Kimura and Onami, 2010; Kimura and Kimura, 2011b; Howard and Garzon-Coral, 2017). These forces move the centrosomes to drive translational and rotational movements of the mitotic spindle. The position and orientation of the mitotic spindle is critical for the size asymmetry and direction of cell division (Morin and Bellaïche, 2011; Siller and Doe, 2009; Gönczy, 2008).

The mechanics controlling the configuration (i.e., position and orientation) of the bipolar spindle is well studied. In contrast, little is known about the mechanics controlling the configuration of the mitotic spindle with 3 or more poles (i.e., a multipolar spindle). Multipolar spindles are formed when cells possess more than 2 centrosomes owing to a defect in the regulation of their numbers (Godinho and Pellman, 2014; Pihan et al., 1998). The forces controlling configuration should be similar for both multipolar and bipolar spindles. Therefore, multipolar spindles might provide a good example to test the feasibility of the theories proposed for the regulation of bipolar spindles. In addition, the configuration of multipolar spindles might be important to understand the viability of cancer cells. Supernumerary centrosomes are frequently observed in cancer cells and are expected to induce multipolar spindles, and thus aneuploidy and cell death (Boveri, 2008; Lingle et al., 1998; Brinkley, 2001; Holland and Cleveland, 2009). However, cancer cells are known to proliferate efficiently, which presents a paradox (Godinho et al., 2009). One mechanism to overcome the paradox is to cluster supernumerary centrosomes into two to form a bipolar spindle (Kwon et al., 2008). However, little is known how spindles behave once multipolar spindles are formed.

In this study, we investigated the configuration of tripolar spindles and consecutive cell division pattern in *Caenorhabditis elegans* embryos. The configuration of bipolar spindles is well established in *C. elegans* embryos (Gönczy and Rose, 2005), and hence it a good system to analyze the configuration of tripolar spindles. To induce reproducibly tripolar spindles in *C. elegans* embryos, we focused on an *emb-27^APC6^* mutant. *C. elegans emb-27* encodes a subunit of anaphase-promoting complex (APC) that is required for the initiation of chromosome segregation and other events at anaphase (Golden et al., 2000). Sperms from *emb-27* mutants do not contain chromosomes, but can fertilize eggs (Sadler and Shakes, 2000). After fertilization, some embryos divide into 3 cells by forming 2 cytokinetic furrows at the first cell division, possibly by forming tripolar spindles (Sadler and Shakes, 2000). In this manuscript, we call the cytokinesis that forms 2 cytokinetic furrows and divides the cell into 3 as “2-furrow cytokinesis”, whereas “1-furrow cytokinesis” refers to a usual cytokinesis with 1 cytokinetic furrow that divides the cell into 2. We have recently shown that the paternal *emb-27* mutant embryos possesses 3 or more centrosomes (Kondo and Kimura, 2018), as expected from the previous report (Sadler and Shakes, 2000). An unexpected result was the frequency of cells with 3 or more centrosomes in the mutant embryos was about 70% (Kondo and Kimura, 2018). This high frequency is seemingly inconsistent with the defective mitosis observed only in one-third of the embryos (Sadler and Shakes, 2000). In this study, we investigated the mechanism how some cells with 3 centrosomes avoid 2-furrow cytokinesis in paternal *emb-27* mutant embryo. This investigation should give an insight into how centrosomes (spindle poles) behave in normal and abnormal conditions.

## Results

### Abnormal number of centrosomes does not always cause the excess number of furrows

We have previously quantified the number of the centrosomes in paternal *emb-27* mutant embryos and found that ∼70% of the mutant embryos possessed 3 or more centrosomes (Kondo and Kimura, 2018). This seemed inconsistent with the previous report that only one-third of the mutant embryos showed defective mitosis (Sadler and Shakes, 2000). To investigate the relationship between the excess number of centrosomes and the mitosis defect, we quantified the number of cell-division furrow in the paternal *emb-27* mutant embryos. About one-third of the paternal *emb-27* embryos at one-cell stage formed 2 cell-division furrows and divided into 3 cells (“2-furrow cytokinesis”, Fig. 1A). The result was in agreement with the ratio of the previous study (Sadler and Shakes, 2000). Only about 30% of the cells possessing 3 centrosomes underwent 2-furrow cytokinesis (Fig. 1B). Moreover, about 20% of the cells with 4 centrosomes still underwent 1-furrow cytokinesis. We did not observe 3-furrow cytokinesis for the cells with 4 centrosomes during the course of this study (Figs. 1B). Therefore, the excess number of centrosomes does not always induce multipolar mitosis.

**Figure 1.**
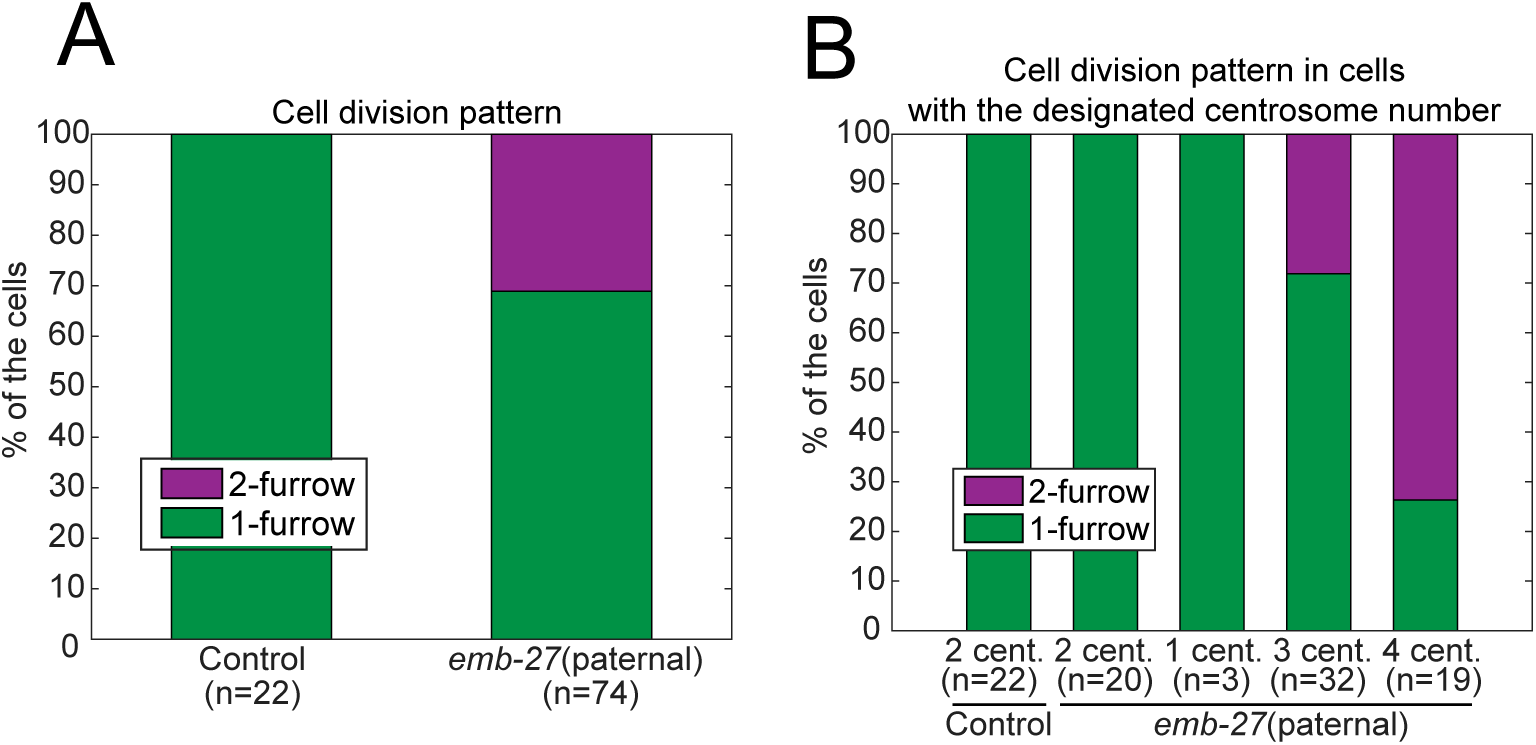
The number of centrosomes and furrows in the paternal *emb-27* mutant embryos. (A) Frequency of two patterns of the first cell division in control and *emb-27* paternal embryos. (B) Frequency of two patterns of the first cell division with the designated number of centrosomes.

### Only 2 of the 3 sides of the tripolar spindle is occupied by chromosomes

To understand the mechanism that determines the choice between 1-furrow and 2-furrow cytokinesis, we focused on the 1-cell stage embryo with 3 centrosomes. Observations of centrosomes and chromosomes in these cells revealed that cells with 3 centrosomes always formed a tripolar spindle, instead of multiple centrosomes merging to form a bipolar spindle (Ring et al., 1982; Quintyne et al., 2005). Interestingly, we noted an unexpected feature that the chromosomes resided only on 2 of the 3 sides of the tripolar spindle in every cell (*n* = 32, Fig. 2AB). Chromatids residing in 2 of the 3 sides of a tripolar spindle have been observed in other cell types (Wilson, 1925; Wheatley and Wang, 1996; Eckley et al., 1997); however, to our knowledge, this is the first report that such a spindle is always observed. We could not determine the mechanism for this event. Nonetheless, this feature on chromatid positioning could be important to understand the difference between 1-furrow and 2-furrow cytokinesis since chromatids are considered critical for the formation of cleavage furrows (Margolis and Andreassen, 1993; Wheatley and Wang, 1996; Eckley et al., 1997).

**Figure 2.**
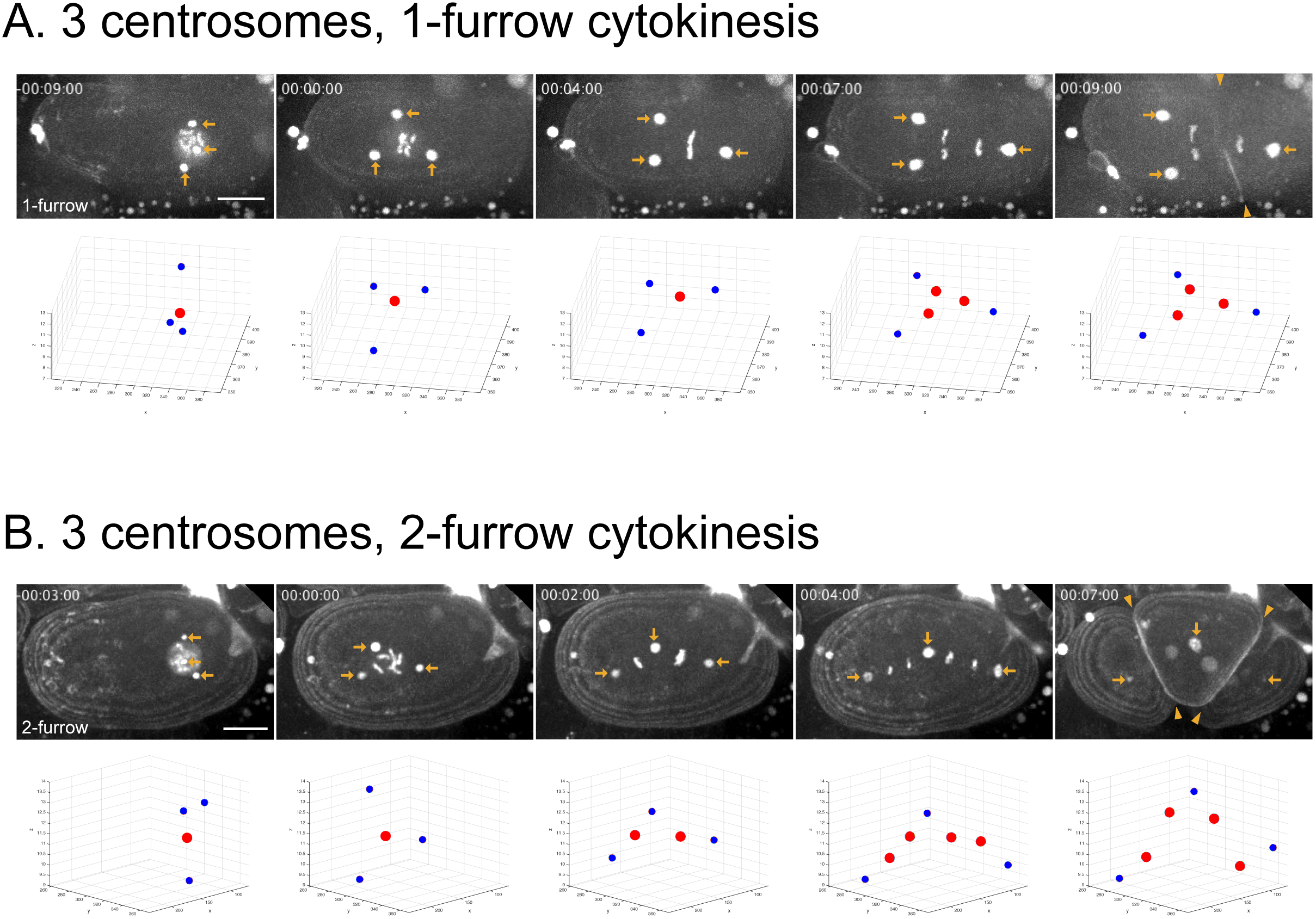
1-furrow cytokinesis and 2-furrow cytokinesis in embryos with 3 centrosomes. Representative time-lapse images of an embryo with 3 centrosomes during the first cell division. Upper panels show the embryos that expressed GFP-tagged γ-tubulin (centrosome, arrows), PH^PLCδ1^ (cell membrane), and histone H2B (nucleus) *in utero*. Lower planes show the position of centrosomes (blue) and nucleus/chromatids (red). In 1-furrow cytokinesis (A), a cleavage furrow (arrowheads) was observed between separated chromatids, which are similar to normal embryos. In 2-furrow cytokinesis (B), two cleavage furrows (arrowheads) were observed. Both patterns have in common that the chromatids are localized only on 2 of the 3 sides of the tripolar spindle. Times are with respect to NEBD. Bar, 10 µm.

### Cell geometry determines the furrowing patterns in embryos with 3 centrosomes

We compared the cells undergoing 1-furrow (Fig. 2A) and 2-furrow cytokinesis (Fig. 2B) and found a difference in the configuration of tripolar spindles at metaphase. To characterize the configuration of tripolar spindles, we focused on the angle of the side without chromatids (“non-chromosome side”) against the long axis of a cell (i.e., the anterior-posterior axis; Fig. 3A). When the angle of the non-chromosome side was near −90°, a cell tends to undergo 1-furrow cytokinesis, whereas the non-chromosome side was near 0° when a cell underwent 2-furrow cytokinesis (Fig. 3B). After spindle elongation in late metaphase and anaphase, only the chromosome sides appeared to elongate actively (Figs. 2A, 2B, 3C) and preferred to align along the AP axis possibly because of the availability in space. In contrast, the non-chromosome side did not elongate considerably in a cell undergoing 1-furrow cytokinesis (Figs. 2A, 3C-i, 3D-i), or elongated passively because of the active elongation of the chromosome sides in a cell undergoing 2-furrow cytokinesis (Figs. 2B, 3C-ii, 3D-ii).

**Figure 3.**
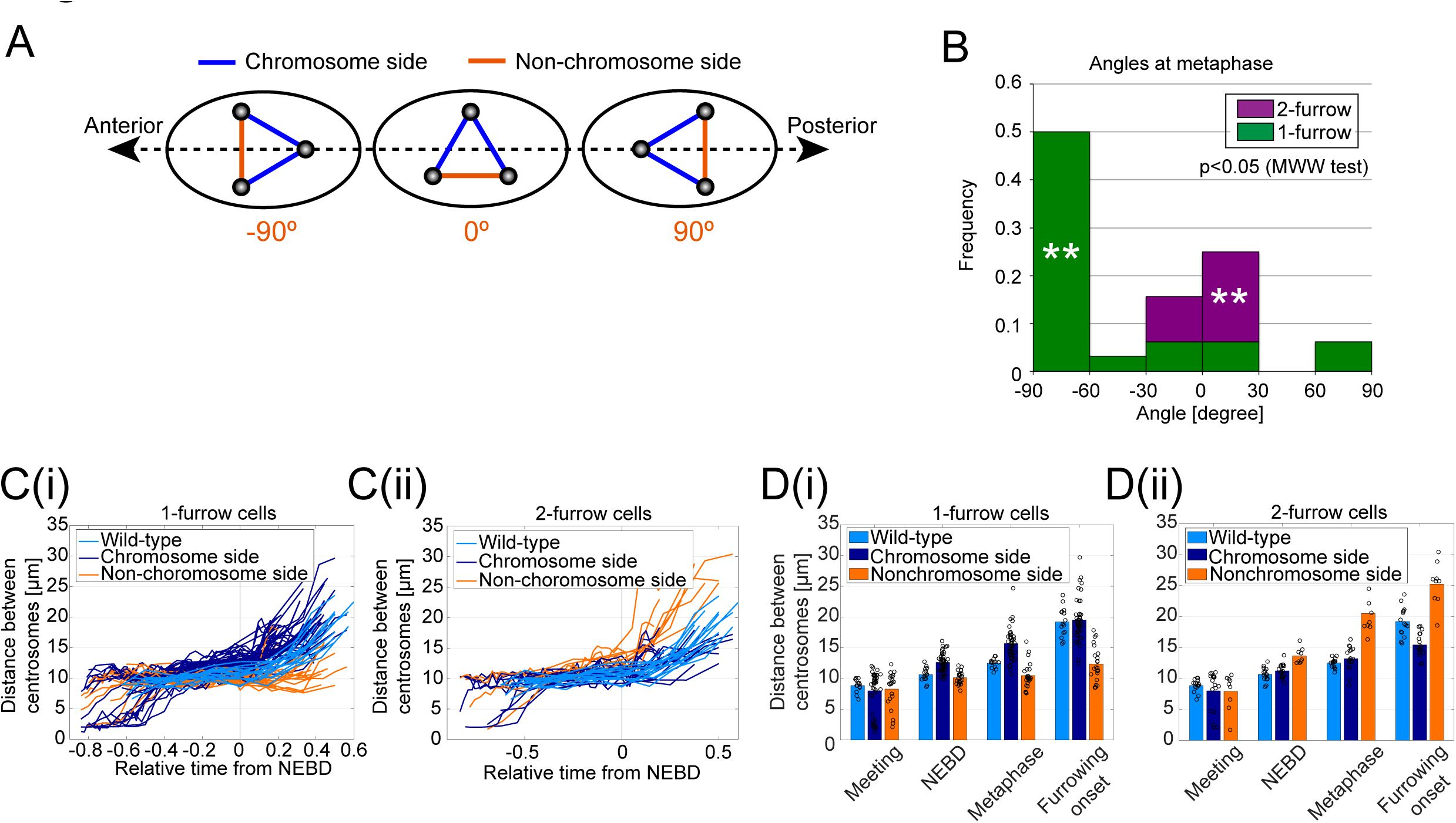
Characterization of tripolar spindles in embryos with 3 centrosomes. (A) Definition of the angle of the side without chromatid (non-chromosome side) with respect to the anterior-posterior axis. (B) Frequency of 1-furrow/2-furrow cytokinesis in embryos with the designated angle of non-chromosome side at metaphase. *n* = 32. **: *p* < 0.005, binomial test for each angle range. (C) Length of the designated side of tripolar spindle over time. Each line represents data from one side. *n* = 32. Time (*T*) is normalized as follows: *T*_relative_ = (*T* -*T*_NEBD_)/([*T*_furrowing onset_ - *T*_NEBD_] - [*T*_pronuclear meeting_ - *T*_NEBD_]). (D) Length of the designated side of tripolar spindle at the designated cellular event. Bars represent mean of all data shown by circles.

From the above observations, we propose the mechanism underlying the difference between 1-furrow and 2-furrow cytokinesis as follows with 2 assumptions (Fig. 4A). [Assumption 1] At anaphase, all 3 poles are pulled outward because the pulling forces act on astral microtubules, like the usual elongation of bipolar spindles (Grill et al., 2001; Hara and Kimura, 2009). The directions of forces are geometrically dependent: stronger along the AP-axis like in the elongation of bipolar spindles, possibly because the pulling forces are stronger for longer microtubules (Hamaguchi and Hiramoto, 1986; Kimura and Onami, 2005; Minc et al., 2011). [Assumption 2] Among the 3 sides of the tripolar spindle, 2 chromosome sides resist against the elongation, whereas the non-chromosome side does not. An intuitive consequence of the assumptions is that, for tripolar spindles with near 0° configuration, the angle between the chromosome sides might increase, and the cells will form 2 furrows; in contrast, for spindles near with −90° or +90° configuration, the angle might decrease, and the cells will form 1 furrow.

**Figure 4.**
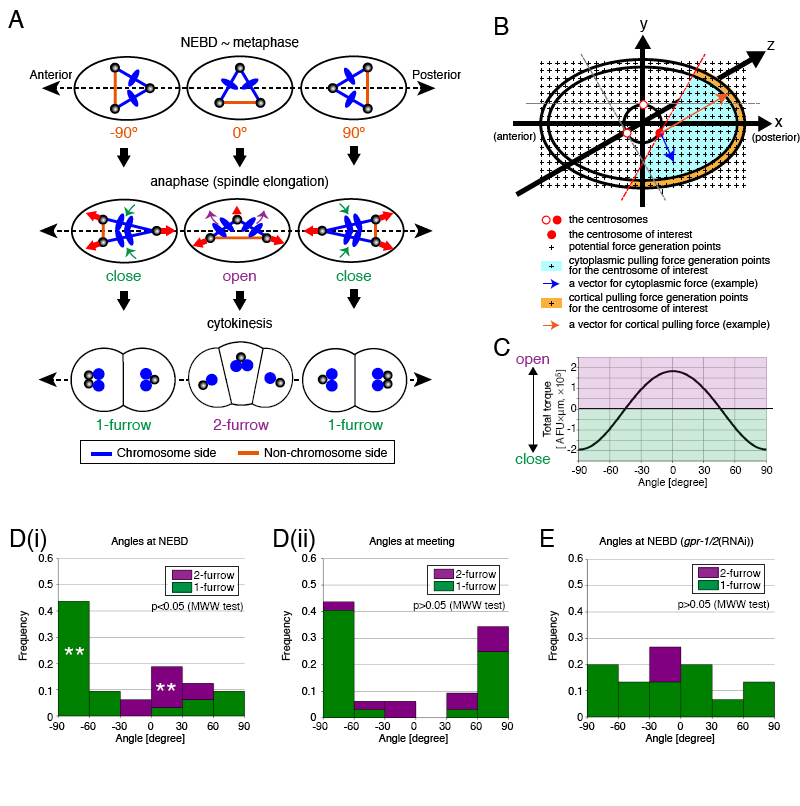
A numerical model to relate the angle of the tripolar spindle and the decision between 1-furrow and 2-furrow cytokinesis. (A) A scheme of the proposed model for how 1-furrow and 2-furrow cytokinesis are determined depending on the angle of tripolar spindle. The upper panel is the same as that in Figure 3A, showing the angle of tripolar spindle at NEBD to metaphase. The middle panel shows the elongation of the tripolar spindle. We assume only the chromosome sides (blue) elongate actively, whereas the length of non-chromosome side (orange) depends on the movements of the 2 ends. The forces pulling the pole (centrosome) depend on the ellipsoidal geometry of the cell (red arrows), and the angle between the chromosome sides close (green arrows) or open (purple arrows) depending on the direction of forces. The opening or closing induces 1-furrow or 2-furrow cytokinesis, respectively (the lower panel). For 2-furrow cytokinesis, 2 of the 3 daughter cells inherit only half the set of the chromosomes (blue), resulting in aneuploidy. (B) Our numerical 3D model to calculate the forces acting on each of the 3 poles that pull a pole (e.g., red circle) with a geometry-dependent force and cortical pulling forces (e.g., orange arrow, from force generators at the orange region). As geometry-dependent force, we assumed cytoplasmic pulling forces (e.g., blue arrow, from force generators at the blue region). (C) Calculation of the torque to open or close the angle between the chromosome sides (Fig. 4A, middle panel), depending on the angle of tripolar spindle (Fig. 4A, upper panel). (D) Frequency of 1-furrow or 2-furrow cytokinesis in embryos with the designated angle of non-chromosome side at NEBD (i) and pronuclear meeting (ii). *n* = 32. **: *p* < 0.005, binomial test for each angle range. (E) Frequency of 1-furrow or 2-furrow cytokinesis in *gpr-1/2* (RNAi) embryos with the designated angle of non-chromosome side at NEBD. *n* = 15.

This proposal was tested quantitatively by developing a 3D force calculation model. As the initial condition, we placed an equilateral triangle, which corresponds to the tripolar spindle, at the center of a prolate ellipsoid, corresponding to the cell. The 3 vertices of the triangle were pulled outward depending on the geometry of the cell (Fig. 4B, and Methods). The torque to open or close the angle between the 2 chromosome sides was calculated under the condition that the non-chromosome side does not resist against elongation or compression forces. The calculation confirmed the intuitive consideration that tripolar spindles with near 0° configuration tend to “open”, whereas those with near −90° or +90° configuration tend to “close”, leading to the induction of 2-furrow and 1-furrow cytokinesis, respectively (Fig. 4C). Therefore, the angle of tripolar spindles and the geometry-dependent forces on the poles account for the choice between 1-furrow and 2-furrow cytokinesis.

### The mechanisms controlling the initial angle of tripolar spindles: Experimental observations

Since we proposed above that the angle of tripolar spindles against the long axis of a cell determines the choice between 1-furrrow and 2-furrow cytokinesis, we next investigated the mechanism to determine the initial angle of the spindle before elongation. We first determined when this angle critical for the choice is established. The asymmetry in the length of the 3 sides of tripolar spindles was not evident at nuclear envelope breakdown (NEBD; Fig. 3D). In contrast, the angle of the future non-chromosome side was already biased between 1-furrow and 2-furrow cytokinesis at NEBD, but not at the earlier stage of pronuclear meeting (Fig. 4D-i, 4D-ii, 2-furrow (purple) vs. 1-furrow (green)). At pronuclear meeting, the frequency of observing 1-furrow or 2-furrow cytokinesis at each angle range was not statistically biased (*p* > 0.05) compared to the overall frequency of observing 1-furrow or 2-furrow cytokinesis. We propose that the angle of the triangle made by the 3 centrosomes at NEBD, but not earlier (i.e., meeting), is critical for the choice between 1-furrow and 2-furrow cytokinesis.

We also found that the angle distribution of tripolar spindles at NEBD was not symmetric against the AP axis (Figs. 3B, 4D-i). The −90° configuration was more favorable than the +90° configuration. In the *C. elegans* embryos, the centrosomes are known to be pulled by the force generators residing at the cortex via the microtubules (Grill et al., 2001). The strength of the cortical pulling force is asymmetric along the AP axis, which results in asymmetric positioning of the spindle and cell division plane. The cortical pulling forces are dependent on *gpr-1/2* gene products (Srinivasan et al., 2003; Colombo et al., 2003). If the asymmetry of the angle of tripolar spindles is regulated by the asymmetric cortical pulling force, such asymmetry should be lost by the knock-down of *gpr-1/2*. In fact, this was the case (Fig. 4E). Therefore, the cortical pulling forces affect the angle of tripolar spindles to generate its asymmetric distribution along the AP axis.

### A numerical model to predict favorable angles of tripolar spindles at NEBD

We next asked whether we can construct a numerical model to account for the favorable angles for the tripolar spindles at NEBD (experimental observations: Fig. 4D-i, 4E). For this purpose, we use the 3D force calculation model constructed to calculate the opening or closing of the tripolar spindles at anaphase (Fig. 4B). By using this framework, the favorable angle was predicted based on a method developed previously to calculate the orientation of bipolar spindles in mammalian cultured cells (Théry et al., 2007; Matsumura et al., 2016). When only the ellipsoidal geometry of a cell was considered, the distribution of the angle of the non-chromosome side was uniform (Fig. 5A), which was consistent with the experimental observation for *gpr-1/2* (RNAi) condition that is defective for asymmetric cortical pulling force (Fig. 4E).

**Figure 5.**
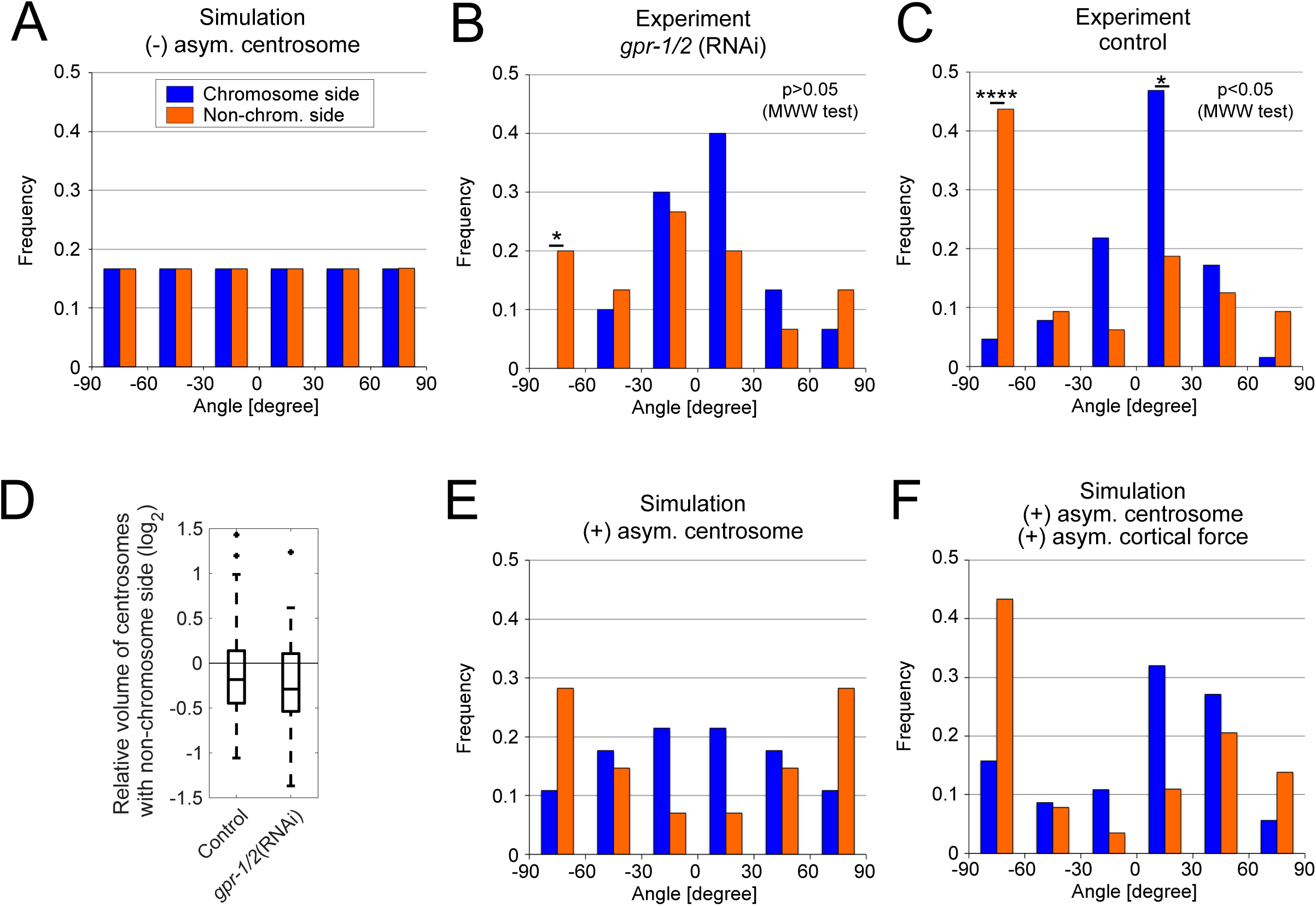
The heterogeneity in centrosome size accounts for the different distributions between the angles of chromosome and non-chromosome sides. (A) The simulated angle of the sides of tripolar spindle before elongation. The angle of the side is 0° when the side is parallel to the long axis and +90° or −90° when perpendicular (see the orange side in Figures 3A or 4A), for both chromosome (blue) and non-chromosome (orange) sides. This definition is common throughout. (B, C) The experimental distribution in *gpr-1/2* (RNAi) (B) and control (C) at NEBD. *n* = 45 for *gpr-1/2* (RNAi) and *n* = 96 for control. *: *p* < 0.05, ****: *p* < 5 × 10^-5^ binomial test for each angle range. (D) The experimental heterogeneity in centrosome size at NEBD. The average volume of the 2 centrosomes of the ends of the non-chromosome side was divided by the volume of the centrosomes between the chromosome sides. The logarithmic ratio is shown with a box plot drawn using MATLAB software. The ratios are smaller than 1 in all conditions examined. (E-F) The simulated distribution of the angle in a model with heterogeneity in centrosome size (E), which should reflect the experimental condition of *gpr-1/2* (RNAi) (B), and after adding the asymmetric cortical pulling forces (F), which should reflect the control experiment (C).

This model condition was axially symmetric with respect to the long axis of the cell, and thus did not discriminate between the chromosome and non-chromosome sides (Fig. 5A). In contrast, *in vivo*, the non-chromosome sides tend to position near +90° or −90°, whereas the chromosome sides tend to position near 0° for *gpr-1/2* (RNAi) (Fig. 5B). The difference in the angle distribution of non-chromosome and chromosome sides was statistically significant for control embryos (Fig. 5C, *p* < 0.05 as per Mardia-Watson-Wheeler test). The disagreement between the model and the experiments indicated that the non-chromosome side needs to be determined non-randomly through an uncharacterized mechanism.

### Heterogeneity in centrosome size explains the angle of tripolar spindles

Our observation that chromosomes reside on 2 out of 3 sides of a tripolar spindle implies that the 3 poles (centrosomes) of the spindle are not equivalent. Among the 3 poles, only 1 is connected to 2 sides both with the chromosome, whereas the remaining 2 poles are connected to the chromosome and non-chromosome side each. We call the former as “chromosome poles”, whereas the latter as “non-chromosome poles”. We found that the chromosome poles tended to be larger in size than the non-chromosome ones at NEBD (Fig. 5D). In binary comparison, where a chromosome pole is compared for each of the non-chromosome poles, the chromosome poles were larger with statistically significant frequency [*n* = 43/64 (*p* < 10^-7^) for control, and *n* = 33/46 (*p* < 10^-6^) for *gpr-1/2* (RNAi)]. The chromosome pole was not always the largest among the three poles, but it was rarely the smallest [*n* = 1/32 (*p* < 10^-4^) for control, and *n* = 1/23 (*p* < 10^-2^) for *gpr-1/2* (RNAi)]. Because the measured sizes of the centrosomes depend on their relative position to the focal planes of microscopy (2 μm intervals in this measurement), the measurement of size might involve some experimental errors. Considering that the abovementioned statistically significant tendency of the chromosome pole to be large, and the possible experimental error, we concluded that one of the large poles (centrosome), possibly the largest, is selected to become the chromosome pole.

Large centrosomes have been reported to be associated with more number of microtubules (Greenan et al., 2010). Therefore, we modified our numerical simulation by assuming the number of microtubules growing from the chromosome pole to be larger than that growing from the other 2 poles. As the magnitude of the asymmetry was obscure, we searched for parameters that allowed the model to fit with the experimental results. This simulation reproduced the alignment of the tripolar spindle *in vivo*, in which the non-chromosome side tends to assume +90° or −90° configuration, whereas the chromosome sides tend to have 0° configuration (Fig. 5B–experiment, 5E–simulation). The simulation result, together with the experimental observation, indicates that heterogeneity in the size of the centrosomes exists, and the largest centrosome captures chromosomes at the 2 sides, whereas the other centrosomes capture chromosomes only at 1 side. This heterogeneity in centrosome size accounts for the alignment of the tripolar spindle.

When cortical pulling forces were added to the simulation (Kimura and Onami, 2007; 2010), we could further reproduce the asymmetry along the AP axis of the alignment of the tripolar spindle (Fig. 5C–experiment, 5F– simulation). This allowed the non-chromosome side to be concentrated near −90°, but not at +90°.

The results collectively support the model, in which the arrangement (i.e., the angle with respect to the long axis) of the tripolar spindle is determined by 3 factors: the forces to pull the spindle poles depending on the cell geometry, heterogeneity of the centrosome size, and asymmetric cortical pulling forces (Fig. 4A, 4B). This angle of the tripolar spindle determines whether a cell undergoes 1-furrow or 2-furrow cytokinesis, which leads to the inheritance of normal chromosome number or aneuploidy, respectively. The determination of the spindle arrangement depending on the 3 factors should not be limited to the tripolar spindle, but would be generally applicable to all types of spindles.

## Discussion

In this study, we investigated the patterns of cytokinesis induced after the formation of a tripolar spindle. To induce tripolar spindles in the *C. elegans* one-cell stage embryonic cell, we used a paternal *emb-27* (*g48ts*) mutation. This enabled the determination of the effect of multipolar spindles in a cell by using maternal, wild-type *EMB-27* gene product. Therefore, the phenotypes observed for the cells were determined to be the consequence of the defects in sperm. Notably, the *emb-27* sperm not only has multiple centrosomes, but is also anucleated (Sadler and Shakes, 2000). The resultant zygotes are thus haploid, and we cannot exclude the possibility that the ploidy affected the angle of the spindle and the pattern of cytokinesis. Although we think that this scenario is unlikely, to exclude the possibility, observing *emb-27* mutant sperms with multiple centrosomes but not anucleated, which are obtained with a quite low frequency, will be informative.

Since excess number of centrosomes was delivered into a zygote with an *emb-27* sperm, the mitotic spindle formed in the one-cell stage embryonic cell became multipolar (Fig. 1). In this study, we focused on the tripolar spindle, because it is the simplest form of multipolar (> 2 poles) spindles. When a tripolar spindle forms, it does not always divide into 3 cells with each daughter cell possessing one centrosome (i.e., 2-furrow cytokinesis), but division into 2 cells (i.e., 1-furrow cytokinesis) was more frequent. With 1-furrow cytokinesis, a daughter cell contains excess number of centrosomes, but the chromosomes are equally segregated (Fig. 4A). Therefore, inducing 1-furrow cytokinesis in a cell with tripolar spindle can be regarded as a mechanism to avoid aneuploidy. The decision whether a cell adopts 1-furrow or 2-furrow cytokinesis was likely dependent on the angle of the tripolar spindle against the long axis of the ellipsoidal cell (Fig. 3AB). Unexpectedly, in the tripolar spindle of this study, only 2 of 3 sides were occupied by chromosomes. The 2 sides correlated with the position of the large centrosome (Fig. 5D). However, considering a mechanism in which some quantitative changes in the centrosome size result in the occupation of the chromosomes in all-or-none manner is difficult. A novel mechanism might exist to regulate the occupation of chromosomes in a mitotic spindle, which needs to be investigated in the future.

We found that the major determinant of the decision between 1-furrow and 2-furrow cytokinesis is the angle of the tripolar spindle at NEBD: if it is around 0°, the cell will undergo 2-furrow cytokinesis; if it is around −90° or +90°, a cell will undergo 1-furrow cytokinesis (Fig. 3). The link between the angle and the patterns of cytokinesis was explained numerically by assuming the force to pull the poles of the spindle in a geometrical manner (Fig. 4). Furthermore, we succeeded to explain the tendency in the distribution of the angle at NEBD by using the same theoretical framework by adding asymmetric cortical pulling forces and considering the heterogeneity in the size of the 3 centrosomes (Fig. 5). In conclusion, we propose that the angle of the tripolar spindle and the patterns of cytokinesis are regulated by 3 factors: the cell-geometry-dependent forces, cell-polarity-dependent cortical pulling forces, and the heterogeneity in centrosome size. These mechanisms suggest why a majority of cells with tripolar spindles avoid aneuploidy in the one-cell *C. elegans* embryos.

Applying this mechanism directly to other cells/organisms might be difficult. The *C. elegans* embryonic cell is unique in terms of its ellipsoidal shape and the feature where only 2 of the 3 sides of the tripolar spindle are occupied by chromatids. However, our study results clearly indicate the possible contribution of cell geometry to the behavior of multipolar mitotic spindles, and thus might provide a new insight when considering abnormal cell division induced by excess number of centrosomes, such as in the case of cancer. Further, the present study characterized the forces involved in the positioning of normal, bipolar spindle, which are important for symmetric and asymmetric cell divisions.

## Materials and methods

### Worm strains and maintenance

The strains used in this study are listed in Table S1, and made in our another study (Kondo and Kimura, 2018). Strains were maintained under standard conditions (Brenner, 1974). To obtain paternal *emb-27* mutant embryos, temperature shift and mating were conducted as follows: 10 gravid hermaphrodites of CAL0051 were placed on a fresh 60 mm plate and allowed to lay eggs at 16ºC. After 21-28 h, the adults were removed from the plate, and the plate was incubated for another 26-30 h at 16ºC. Next, the plate was transferred to 25ºC and incubated for another 13-18 h. Subsequently, 30 males from the plate were moved to a new 35 mm plate with 10 hermaphrodites of CAL0182 (*fem-1*ts), which were grown at 25ºC from L1/L2 stage to prevent self-fertilization, for 13-18 h to induce mating, and the embryos from the hermaphrodites were observed.

### Live cell imaging microscopy: Observation of embryos

For observation of embryos *in utero*, anesthetized adult worms were placed on 2% agar pad and gently sealed with a coverslip (Kimura and Kimura, 2012). The samples were observed using a spinning-disk confocal system (CSU-X1; Yokogawa Electric, Tokyo, Japan) mounted on a microscope (IX71; Olympus). Images were acquired every minute (for *in utero*) for the thickness of 30 µm with 2 μm z-intervals at 20 ms exposure by using a UPlanSApo 60× 1.3 NA objective (Olympus) equipped with an EM-CCD camera (iXon; Andor, Belfast, UK) controlled by Metamorph software (Molecular Devices, Sunnyvale, CA, USA).

### Image processing and analysis

For the measurement of distance *L* between centrosomes in Figure 3, the coordinates of each centrosome in 3D (*x*, *y*, and *z*) were obtained manually from the images by using ImageJ, and the Euclidean distance was calculated. For the computation of angle in Figures 3-5, the coordinates of centrosomes and AP poles in 3D were measured manually from the images, and the angle against the AP axis was calculated using a custom-written code in MATLAB. In control cells, the angle *θ_wt_* was calculated using the formula: *θ*_wt_ = abs(-[acos{dot(**BI**, **AP**)/norm(**BI**) × norm(**AP**)} × 180/π] + 90), where **BI** and **AP** are the unit vectors connecting the two poles of the bipolar spindle and the unit vector of AP axis, respectively. In the *emb-27* mutant embryos with a tripolar spindle, the angle of non-chromosomal side to the AP axis *θ_tri_* was calculated using the formula: *θ_tri_* = acos{dot(**TRI**, **AP**)/norm(**TRI**) × norm(**AP**)} × 180/π, where **TRI** is the unit vector of the non-chromosome side. If *θ* ≥ 90º, the value was subtracted from 180º and replaced with the original *θ.*

Measurement of centrosome size using γ-tubulin::GFP fluorescence (Fig. 5D) was conducted as described in our another study (Kondo and Kimura, 2018).

### Numerical simulation to calculate the torque to open or close the tripolar spindle at anaphase (Fig. 4C) or to calculate stable angles of the tripolar spindle at NEBD (Fig. 5)

We constructed a 3D simulation assuming the cell as an ellipsoid, with one major axis of 25 μm in radius and two minor axes of 15 μm in radius, which is similar to the actual cell (Fig. 4B, Table S2). Since we focused on the angle of the entire tripolar spindle (Fig. 5) or the angle of the chromosome sides (Fig. 4C), we fixed the center of the spindle at the center of the cell and only considered rotational movements around the center of the spindle (Fig. 5) or the center of the chromosome sides (Fig. 4C). We also assumed the tripolar spindle to be an equilateral triangle inscribed in a circle with the radius of 5 μm on a plane (x-y plane in Fig. 4B), including the AP axis, which is similar to the situation from NEBD to metaphase in the actual cell when the rotation occurs.

#### Forces acting on the spindle poles

We assumed two kinds of forces to act on the spindle pole (Kimura and Onami, 2007; 2010). One is the geometry (cell shape)-dependent force. As for this force, we introduced cytoplasmic pulling force, which pulls the astral microtubule growing from the pole toward outside of the spindle at the cytoplasm (Hamaguchi and Hiramoto, 1986; Kimura and Onami, 2005; Wühr et al., 2010; Kimura and Kimura, 2011a; Tanimoto et al., 2016). In this study, we assumed that the astral microtubules from one pole cover the entire region of the cytoplasm opposite to the spindle (Fig. 4B, light blue region). The direction and magnitude of the cytoplasmic pulling force acting on the pole were calculated by summing up all the unit vectors heading from the pole to each of the “force generation points” evenly distributed in the cytoplasm as the tetragonal lattice points with 0.5 + N(0, 10^-6^) μm interval (Fig. 4B, cross), where N (0, 10^-6^) is a random number following normal distribution with mean at 0 and standard deviation of 0.001 μm to minimize artifacts caused by the regular interval of the lattice. This setting is consistent with the cytoplasmic pulling force proportional to the length of each microtubule (Kimura and Onami, 2005; Minc et al., 2011; Kimura and Kimura, 2011b).

The other kind of force is cortical pulling force, which pulls the poles from the cortex via astral microtubules (Grill et al., 2001). In this study, the direction and magnitude of cortical pulling force acting on the pole were calculated by summing up all the unit vectors heading from the pole to each of the “force generation points” evenly distributed as the tetragonal lattice points in the cortical region (Fig. 4B, crosses in the orange region, which is 2 μm thick). This setting is consistent with the cortical pulling force acting on the pole proportional to the cortical area covered by the astral microtubules (Grill et al., 2003; Grill and Hyman, 2005; Hara and Kimura, 2009; Kimura and Kimura, 2011b). In the present simulation, we set the force generated by one force generation point in the cytoplasm to be 1 arbitrary force unit (AFU), whereas that in the cortical region to be *kf*, and 1.5 × *kf* [AFU] for anterior and posterior cortex, respectively. The *kf* parameter was chosen among {0.2, 0.4, 0.6, 0.8, 1, 2, 3}, and we found *kf* = 2 to yield the best result resembling the experimentally obtained angle distribution (Table S2). The asymmetry in cortical pulling force was assumed to be 1.5-fold stronger at the posterior half based on an experimental estimation (Grill et al., 2003). When we introduced asymmetry in the centrosome size, we increased the forces generated by cytoplasmic force generators pulling the large pole to be *l*-fold. This number was chosen among {1.05, 1.1, 1.15, 1.2, 1.25}, and we found *l* = 1.1 to yield the best result resembling the experimental distribution (Table S2).

#### Calculation of torque to open or close the tripolar spindle at anaphase

For this, we calculated the torque to rotate the 2 chromosome sides individually by summing up all the forces acting on the 2 centrosomes located at the ends of the side. The torque to open the spindle (i.e., increase in the angle between the 2 chromosome sides) was assigned the plus sign, and that to close (decrease the angle) was assigned the minus sign. The torque calculated for each side was summed to yield the total torque in Figure 4C.

#### Calculation of the torque to rotate the tripolar spindle at NEBD

We predicted the stable angle of the tripolar spindle against the long (AP) axis of the embryo by calculating the potential energy landscape as developed by Théry et al. (Théry et al., 2007) and that used in our previous study (Matsumura et al., 2016). By summing up all the forces acting on each pole and summing up the forces acting on the three poles, we calculated the torque acting on the center of the tripolar spindle (Fig. 5). The torque was always acting to rotate the spindle around the z-axis, which was reasonable considering the symmetry in the geometry of the spindle.

#### Calculation of energy and probability

By summing up the torque acting against the attempt to rotate the spindle from 0° configuration (Fig. 4B) to the degree of interest with 1° interval, we calculated the potential energy landscape as described previously (Théry et al., 2007). The unit of the energy (*W*(*θ*)) is [AFU × μm]. The probability to observe a particular angle was calculated as *P(θ) = Nexp(-W(θ)/d)*, where *N* is a normalization factor, and *d* is a coefficient to convert energy to probability (Théry et al., 2007). In this study, *d* was chosen among {10*n* × π/1 80} where *n =* {4, 4.5, 5, 5.5, 6, 6.5, 7}; we found *d =* 5,500 (*n* = 5.5) to yield the best result resembling the experimental distribution (Table S2).

#### Likelihood to explain the experimental results with the simulation

To select the best set of parameters (*kf*, *l*, *d*) in the simulation, we calculated the log-likelihood according to the following equation: [log-likelihood] = Σ*i* = 1^6^log{*C*ns(*i*)P_ns_(*i*)}+ Σ*i*=1^6^log{*C*cs(*i*)*P*cs(*i*)}. Here, *C*ns(*i*) and *C*cs(*i*) are the experimental counts of observing non-chromosome side and chromosome side, respectively, in the *i*-th angle distribution class. The *i*-th angle distribution class is the angle between −90 + 30(*i*-1) to −90 + 30*i* [°]. *P*ns(*i*) and *P*cs(*i*) are the probability in the simulation of observing non-chromosome side and chromosome side, respectively, in the *i*-th angle distribution class.

### Statistical analyses

To test whether the chromosome pole is significantly larger than the non-chromosome poles (Fig. 5D), we conducted binomial test. We tested whether the probability of the chromosome pole being larger than each of the other 2 centrosomes is significantly larger than half. We also tested whether the probability of the chromosome pole being the smallest among the 3 was significantly smaller than one-third.

The statistical difference of angle distributions (Figs. 3B, 4D, 4E, 5B, 5C) was tested in two ways: a binomial test and the Mardia-Watson-Wheeler (MWW) test. In the binomial test, we first calculated the expected probability of observing each group to compare [i.e., 1-furrow (as group A) vs. 2-furrow (as group B), or chromosome side (group A) vs. non-chromosome side (group B)] as *P*A and *P*B (= 1 - *P*A). Next, we calculated the probability of observing group A or B for *n*A or *n*B times or more within *n*A + *n*B trials at the angle range of interest (e.g., −90 to −60°, −60 to −30°) under the assumption that the probability of observing A and B is *P*A and *P*B. The calculation was conducted using Microsoft Excel software.

The MWW test is a non-parametric test to compare the angle distribution of two groups. The calculation was performed both with “hand calculation” by using Excel following the procedure described in Mardia (1967) (Mardia, 1967), as well as with the ‘wason.wheeler.test’ function of R software, and the agreement from both the methods was confirmed. For the comparison between 1-furrow and 2-furrow in *gpr-1/2* (RNAi), we did not use chi-square test, but referred to a table by Mardia (Mardia, 1967) since the sample size was small (*n* = 15).

## Acknowledgments

We thank the members of our laboratory for discussion, Drs. Daiju Kitagawa, Yasushi Hiromi, Mitsuhiko Kurusu, Hiroaki Seino, Kenji Kimura, and Yohei Kikuchi for critical reading of the manuscript, and Drs. Shuji Ishihara (The University of Tokyo) and Shogo Kato (The Institute of Statistical Mathematics) for informative lecture on angular statistics. Some worm strains used in this work were provided by the Caenorhabditis Genetics Center. T.K. was a postdoctoral fellow of the National Institute of Genetics. This work was supported by JSPS KAKENHI (grant numbers: JP15H04372, JP15KT0083, JP16H00816, 18H05529 and 18H02414 to A.K.) and the Naito Foundation and Sumitomo Foundation. The authors declare no competing financial interests.

## Author contributions

T.K. and A.K. conceived the study, constructed the models, executed the calculations, analyzed the data, and wrote the manuscript. T.K. performed all the experiments.

## List of abbreviations

NEBD: nuclear envelope breakdown
AFU: arbitrary force unit

